# Maternal biomarker patterns for metabolism and inflammation in pregnancy are influenced by multiple micronutrient supplementation and associated with childrens’s biomarker patterns and nutritional status at 9-12 years of age in Lombok, Indonesia

**DOI:** 10.1101/625103

**Authors:** Lidwina Priliani, Sukma Oktavianthi, Elizabeth L. Prado, Safarina G. Malik, Anuraj H. Shankar

## Abstract

Maternal nutritional status influences fetal development and long-term risk for adult non-communicable diseases. The underlying mechanisms of these long-term effects remain poorly understood. We examined whether maternal biomarkers for metabolism and inflammation during pregnancy were associated with child biomarkers in the Supplementation with Multiple Micronutrients Intervention Trial (SUMMIT, ISRCTN34151616) in Lombok, Indonesia wherein archived blood specimens and relevant data were available from pregnant women and their children 9-12 years after birth. Forty-four mother-child dyads comprising 132 specimens were analyzed by multiplex microbead immunoassays to quantify vitamin D-binding protein (D), adiponectin (A), retinol-binding protein 4 (R), C-reactive protein (C), and leptin (L). Principal component analysis (PCA) revealed distinct variance patterns, i.e. principal components (PC), for baseline pregnancy bp.pc1.D↓A↓R↓ and bp.pc2.C↓L↑; combined follow-up and post-partum dp-pp.pc1.D↑↓A↑R↑↓L↓ and dp-pp.pc2.A↑C↑L↑; and children ch.pc1.D↑R↑C↑ and ch.pc2.D↓A↑L↑. Maternal multiple micronutrient (MMN) supplementation modified the association between baseline maternal bp.pc2.C↓L↑ and post-supplementation maternal dp-pp.pc2.A↓C↑L↑ (*p*=0.022). Significant associations were found between maternal dp-pp.pc2.A↑C↑L↑ and increased child ‘s ch.pc1.D↑R↑C↑ (*p*=0.036), and decreased child ‘s BMI z-score (BMIZ) (*p*=0.022); and between maternal dp-pp.pc1.D↑↓A↑R↑↓L↑ and increased child ‘s BMIZ (*p*=0.036). Child ‘s ch.pc1.D↑R↑C↑ was associated with decreased birth weight (*p*=0.036), and increased child’s BMIZ (*p*=0.002); and ch.pc2.D↓A↑L↑ was associated with increased child’s BMIZ (*p*=0.005), decreased maternal height (*p*=0.030) and girls (*p*=0.002). Elevated adiponectin and leptin pattern in pregnancy was associated with increased C-reactive protein and vitamin A and D binding proteins pattern in children, suggesting biomarkers acting in concert may be more important than single biomarker effects. Patterns in pregnancy proximal to birth were more associated with child status, and child patterns were most frequently associated with child status, particularly child BMI. Although MMN supplementation and certain maternal biomarker patterns have effects on metabolism and inflammation in pregnancy and in the child, nevertheless, nutrition conditions after birth may have a greater impact.

## INTRODUCTION

Emerging epidemiological evidence has shown that the risk for non-communicable diseases (NCDs) during childhood or as an adult is mediated in part by maternal nutrition in pregnancy and fetal growth [1–3]. Studies in animal models indicate that alterations in nutritional, metabolic, immune and hormonal milieu *in-utero* profoundly affect long-term health of the offspring, including increased risk for NCDs such as diabetes, obesity or cardiovascular disease [4,5]. Knowledge of the underlying mechanisms of these effects remains limited, although evidence is growing for the pivotal roles of metabolism-related hormones and inflammatory mediators [6,7].

Adipocytokines, including leptin, adiponectin and retinol binding protein 4 (RBP4), play an important role in regulating metabolism, energy homoeostasis and inflammatory responses [8–11]. Leptin is involved in body weight control by acting on the satiety center in the hypothalamus [12]. Leptin also acts during pre-natal development by promoting fetal growth and regulating fetal adipose tissue development [13]. Adiponectin plays a role in the catabolism of fatty acids and carbohydrates, improvement of insulin sensitivity and reduction of inflammation [14]. RBP4, previously thought to act as a specific transport protein for retinol, has been added to the family of adipocytokines given its involvement in obesity-induced insulin resistance [15]. Increased concentrations of both leptin and RBP4 have been associated with increased body mass index (BMI) [16,17], while adiponectin concentration was negatively associated with BMI [18]. The concentrations of these adipocytokines during pregnancy have also been associated with adverse pregnancy conditions, including gestational diabetes, preeclampsia and intrauterine growth restriction (IUGR) [19–22]. A previous study reported that maternal leptin and adiponectin concentrations were correlated with fetal leptin and adiponectin concentrations [23].

Inflammatory markers have been associated with increased risk of cardiovascular disease [24]. Specifically, higher C-reactive protein (CRP) concentrations in pregnant women were associated with increased risks for preterm birth and low birth weight (LBW) newborns [25,26], and higher BMI in children [27]. Vitamin D binding protein (VDBP), previously known as a transport protein for vitamin D and as a regulator of vitamin D metabolism [28], has recently been shown to mediate inflammation and macrophage activation [29]. Maternal vitamin D status was reported to have an impact on birth weight and offspring immunity [30,31].

Multiple dietary factors, including micronutrients, have been reported to modulate leptin, adiponectin, RBP4, CRP and VDBP concentrations [32–37]. Maternal expression patterns for these biomarkers may be associated with expression patterns in their children. To examine these relationships, we studied mother-child dyads from the Supplementation with Multiple Micronutrients Intervention Trial (SUMMIT) in Lombok, Indonesia wherein blood specimens and the relevant data were available from pregnancy and children 9-12 years after birth. SUMMIT was a randomized trial comparing maternal multiple micronutrients (MMN) supplementation with iron and folic acid (IFA), and showed that maternal MMN reduced early infant mortality and LBW [38], and noted multiple risk factors for poor fetal development [39]. A follow-up study of children at 9-12 years of age indicated long term effects on child cognitive development. We hypothesized that in this cohort: 1. Maternal nutritional status is associated with maternal biomarkers; 2. Maternal supplementation influenced maternal biomarkers; 3. Maternal biomarkers are associated with child biomarkers; 4. Child biomarkers are associated with child health outcomes (Fig1).

**Fig 1.**
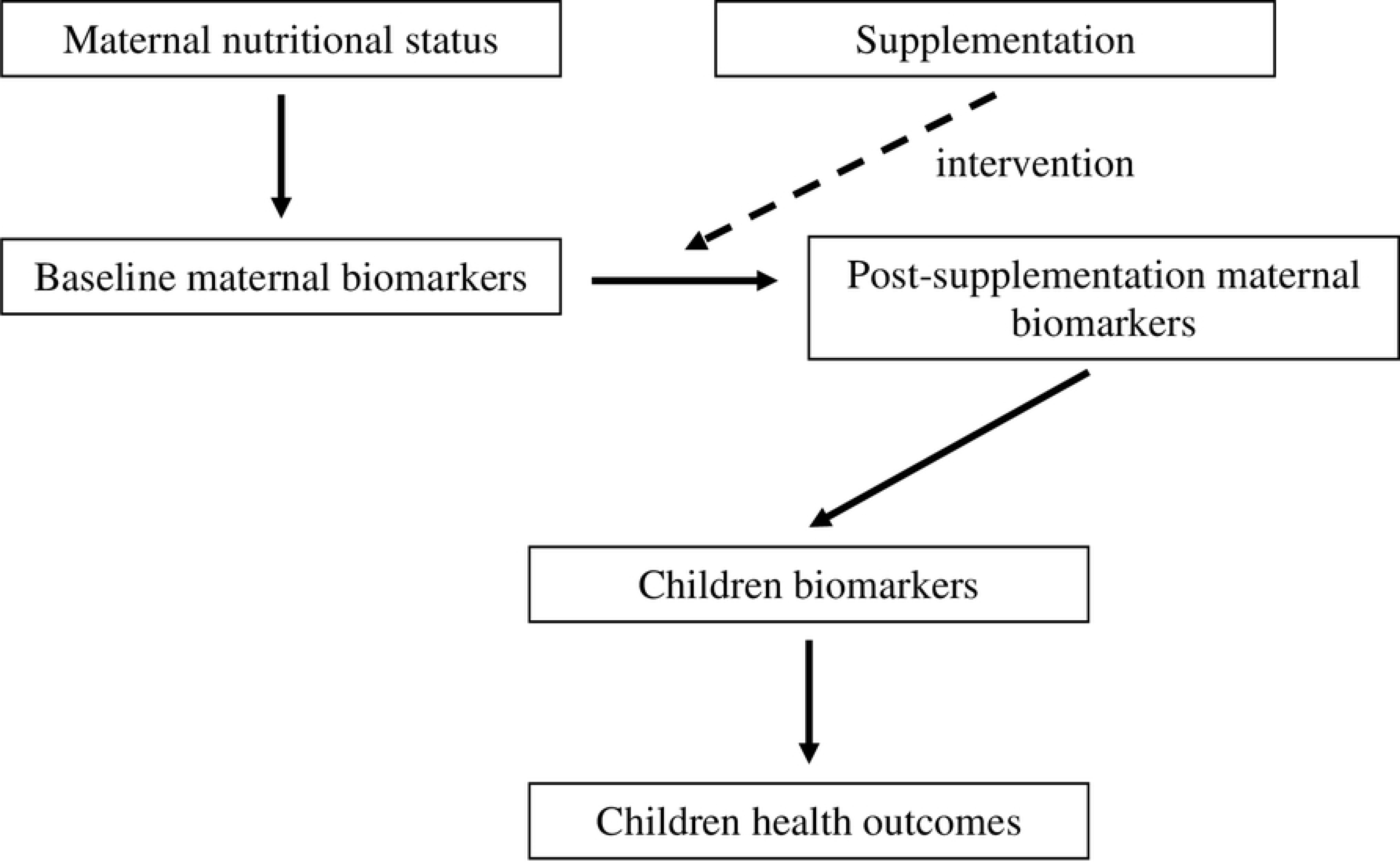
Conceptual framework.

## MATERIALS AND METHODS

### Samples

We selected 414 plasma samples from the SUMMIT mothers and their children. The SUMMIT (ISRCTN34151616) was approved by the National Institute of Health Research and Development of the Ministry of Health of Indonesia, the Provincial Planning Department of Nusa Tenggara Barat Province, and the Johns Hopkins Joint Committee on Clinical Investigation, Baltimore, USA; the follow up study was approved by the University of Mataram Ethical Research Committee as a certified Institutional Review Board of the National Institute of Health Research and Development of the Ministry of Health of Indonesia; the current study of SUMMIT archived materials was also approved by the Eijkman Institute Research Ethics Commission no. 73/2015). Plasma specimens from pregnant women were collected at enrolment before supplementation (baseline) and follow-up specimens at one of four subsequent time points: one month after enrolment, 36 weeks of gestation, one week post-partum, and 12 weeks post-partum (post-supplementation) [40]. From these 414 samples, we further selected 44 maternal plasma at baseline, consisting of 22 samples each of the MMN and the IFA groups, based on a priority list of participants with complete data for other variables collected at multiple time points [40]. Maternal plasma samples from the same participants collected post-supplementation consisted of 18 samples collected during pregnancy (9 samples each of the MMN and the IFA groups), and 26 samples collected postpartum (13 samples each of the MMN and the IFA groups). A total of 44 plasma samples of the children from the aforementioned 44 mothers were selected (Fig 2). Total plasma specimens tested in this study were 132. Maternal nutritional status was characterized by mid-upper arm circumference (MUAC), maternal height and maternal hemoglobin (Hb). Children’s condition was characterized by BMI-for-age z-score (BMIZ), based on World Health Organization norms [41], systolic blood pressure (SBP), and diastolic blood pressure (DBP). Maternal and child biomarkers were determined by VDBP, adiponectin, RBP4, CRP and leptin concentrations.

**Fig 2.**
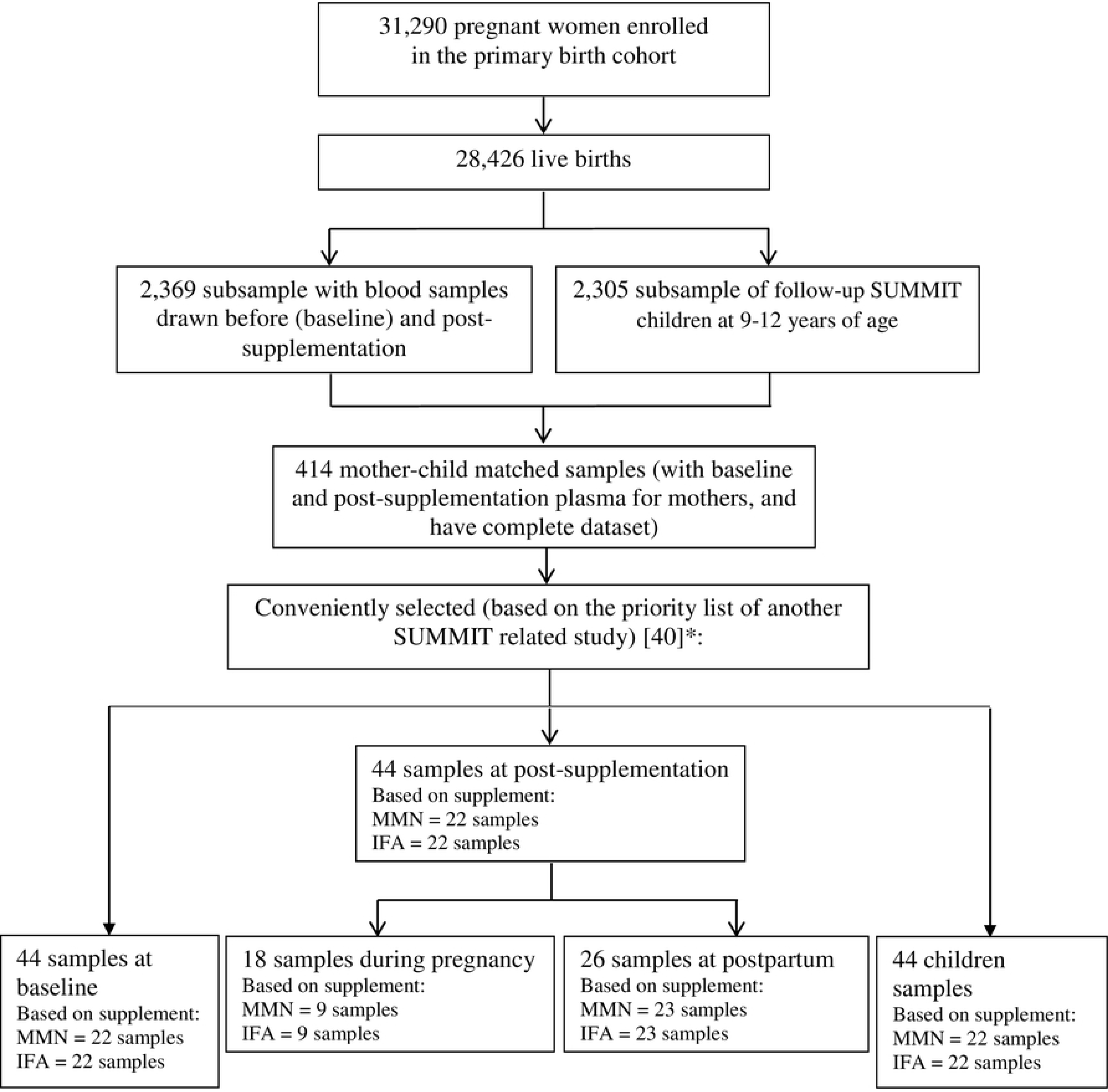
Samples assessment flow chart. IFA = iron folic acid. MMN = multiple micronutrients. 44 paired maternal-children plasma were selected based on a priority list of participants with complete data for other variables collected at multiple time points.

### Multiplex immunoassay

Quantifications of leptin, adiponectin, RBP4, CRP and VDBP were conducted using Luminex^®^ Magnetic Screening Assays (Catalogue number LXSAHM-8, R&D System, Minneapolis, MN, USA) following the manufacturer’s instructions. Briefly, diluted plasma samples were incubated with antibody-coated microspheres, followed by biotinylated detection antibody. Proteins were detected by incubation with phycoerythrin-labeled streptavidin. The bead immuno-complexes were read using MagPix CCD Imager (Luminex, Austin, TX, USA). The instrument was set as follows: events/bead: 50, sample size: 50 μl. Biomarkers concentrations were calculated based on the median fluorescence intensity (MFI), of each duplicate sample.

### Statistical analysis

Data normality was tested by performing the Shapiro Wilk test. Biomarkers concentrations were log-transformed to normalize the data distribution. Normally distributed variables were presented as mean (±standard deviation). Non-normally distributed variables were presented as median (interquartile range). Principal component analysis (PCA) was performed to reduce the five biomarkers into a smaller set of principal components that accounted for most of the variance. We retained the components with eigenvalues of 1 or greater. Factor loadings greater than 0.40 were used to identify biomarkers that loaded on each component, as this threshold value can be used for a structure or pattern coefficient for interpretative purposes. A value of 0.40 would imply that the observed variable shares more than 15% of its variance (0.40^2^ = 0.16) with the component [42].

The principal component (PC) scores from PCA were obtained from each group of samples (baseline, post-supplementation, and child). For multiple linear regression analysis, we combined the post-supplementation PC scores from samples collected during pregnancy and postpartum. Multiple linear regression analysis was performed to determine the associations of (1) maternal PC scores at baseline with maternal nutritional status (association 1), (2) maternal PC scores at baseline with post-supplementation (association 2), (3) maternal PC scores at each time point with child PC scores (association 3), and (4) PC scores of each group of samples with child health outcomes (association 4). Analysis for association 1 (association between maternal PC scores at baseline and maternal nutritional status) employed a regression model with maternal PC at baseline as the outcome and maternal nutritional status such as maternal hemoglobin at baseline, maternal height, and maternal mid-upper arm circumference (MUAC). Analysis for association 2 (association between maternal PC scores at baseline and post-supplementation) employed a regression model with maternal PC at post-supplementation as the outcome and maternal PC at baseline as the predictive variable, adjusting for maternal hemoglobin at baseline, maternal height, maternal mid-upper arm circumference (MUAC), type of supplement (MMN or IFA), baseline to post-supplementation intervals, and timing of post-supplementation (pregnancy or postpartum). We also analyzed the interaction between maternal PC at baseline and type of supplementation with the maternal PC at post-supplementation. The regression models for association 3 (association between children and maternal PC scores) included children’s PC scores as the outcome variable, while maternal PC at baseline and post-supplementation were used as the predictive variables with adjustment for maternal hemoglobin at baseline, maternal height, maternal MUAC, birth weight, child’s gender (boy or girl), type of supplement (MMN or IFA), and timing of post-supplementation (pregnancy or postpartum). Association 4 (children health outcome associations with maternal and children PC scores) was analyzed by regression models which included BMIZ, systolic blood pressure (SBP), and diastolic blood pressure (DBP) of the children as the outcome variables and maternal and child PC scores as the predictive variables. These models included adjustment for maternal hemoglobin at baseline, maternal height, maternal MUAC, birth weight, child’s gender (boy or girl), type of supplement (MMN or IFA), and timing of post-supplementation (pregnancy or postpartum), and an additional adjustment for BMIZ for SBP and DBP regression models. All regression analyses were performed using R-Project for Statistical Computing version 3.4.0, and replicated in some cases with SAS 9.4. The *p*-values of less than 0.05 were considered significant.

## RESULTS

### Baseline characteristics of subjects

The baseline characteristics of mother and child pairs were collected previously during the SUMMIT and its follow up studies, as shown in Table 1. Pregnant mothers who received MMN supplementation had similar characteristics to the ones receiving IFA supplementation. The characteristics of the children whose mothers received MMN or IFA supplementation were similar.

**Table 1.** Baseline characteristics of mother and child pairs.

| Variables | MMN (N = 22) | IFA (N = 22) |
| --- | --- | --- |
| **Mothers** |  |  |
| Age (years) ¶ | 25.0 (20.0-26.5) | 25.5 (20.5-30.0) |
| Parity (number of births) ‡ |  |  |
| 0 | 8 (36) | 5 (23) |
| ≥ 1 | 14 (64) | 17 (77) |
| Height (cm) ¶ | 151.4 (149.3-153.6) | 149.8 (148.7-152.6) |
| Mid-upper arm circumference (mm) ¶ | 239.5 (228.2-253.0) | 245.0 (230.2-253.1) |
| Haemoglobin at enrolment (g/dL) ¶ | 11.1 (10.3-12.0) | 11.3 (10.4-11.9) |
| Gestational age at enrolment (weeks) ¶ | 16.5 (9.5-24.1) | 14.6 (12.3-18.7) |
| **Children** |  |  |
| Gender (M/F) | 13/9 | 10/12 |
| BMI-for-age z-scores † | -0.7 (±1) | -0.8 (±1.1) |
| Systolic blood pressure (mmHg) † | 110.0 (±11.3) | 104.4 (±7.8) |
| Diastolic blood pressure (mmHg) † | 65.0 (±9.8) | 63.4 (±5.3) |
| Birth weight (g) ¶ | 3300 (2925-3500) | 3000 (2825-3450) |
| Gestational age at birth (weeks) ¶ | 39.1 (36.9-40.1) | 39.6 (38.1-40.9) |

¶: median (interquartile range). †: mean (±standard deviation). ‡: n (percentage). MMN: multiple micronutrients supplement. IFA: iron and folic acid supplement.

### Reduction of biomarkers data by Principal Component Analysis (PCA)

From each PCA (maternal baseline, post-supplementation, child), the first two components were retained for further analyses if they had eigenvalues of greater than one. For maternal PCA, percentages of total variance explained by the first two PCs were 60% (PC1 = 39.5%, PC2 = 20.5%) for the baseline group, 77.6% (PC1 = 52.1%, PC2 = 25.5%) for post-supplementation during pregnancy group, and 60.5% (PC1 = 36.9%, PC2 = 23.6%) for post-supplementation postpartum group. For child group, the first two PC explained 63.2% (PC1 = 40.0%, PC2 = 23.2%) of the total variance (Table 2). Each sample group had distinctive components patterns based on biomarker loadings. For the maternal baseline group (bp), PC1 consisted of lower VDBP (D), adiponectin (A) and RBP4 (R), while the PC2 consisted of lower CRP (C) and higher leptin (L), resulting in the variance pattern bp.pc1.D↓A↓R↓ and bp.pc2.C↓L↑. The PC1 of post-supplementation during pregnancy group (dp) was comprised of higher VDBP, adiponectin and RBP4 (dp.pc1.D↑A↑R↑), and PC2 was comprised of higher adiponectin and leptin (dp.pc2.A↑L↑). For post-supplementation postpartum group (pp), PC1 was characterized by lower VDBP, RBP4 and leptin (pp.pc1.D↓R↓L↓), and PC2 by higher adiponectin, CRP and leptin (pp.pc2.A↑C↑L↑). The child (ch) PC1 consisted of higher VDBP, RBP4 and CRP (ch.pc1.D↑R↑C↑), while the PC2 consisted of lower VDBP, and higher adiponectin and leptin (ch.pc2.D↓A↑L↑).

**Table 2.**
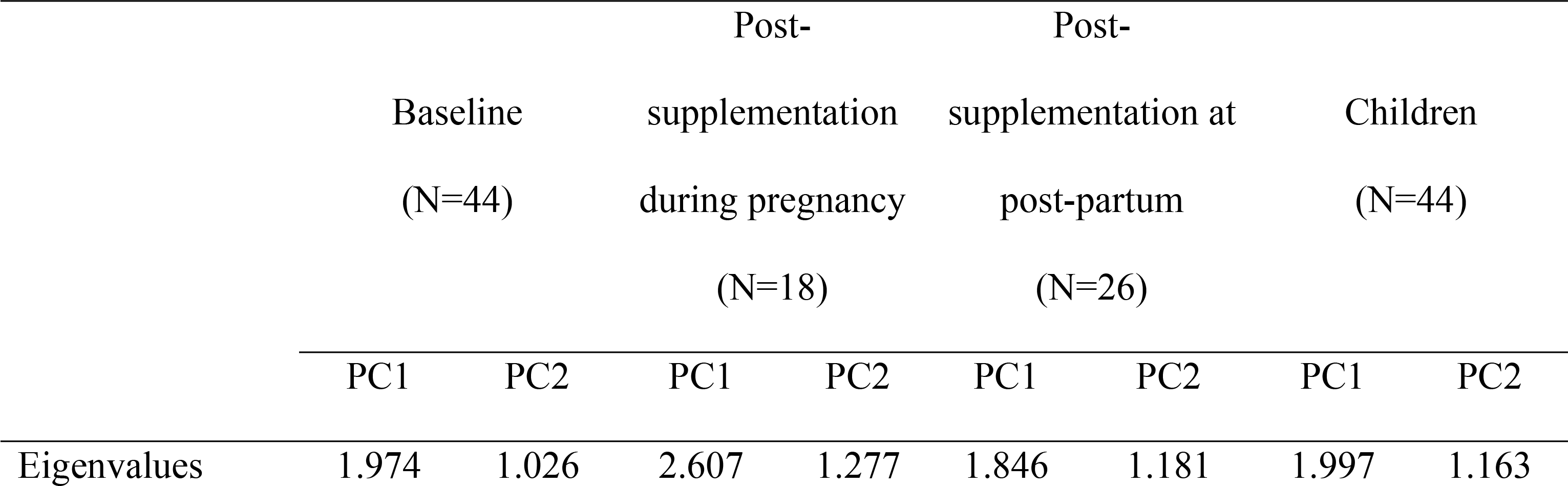
Principal component analysis results.

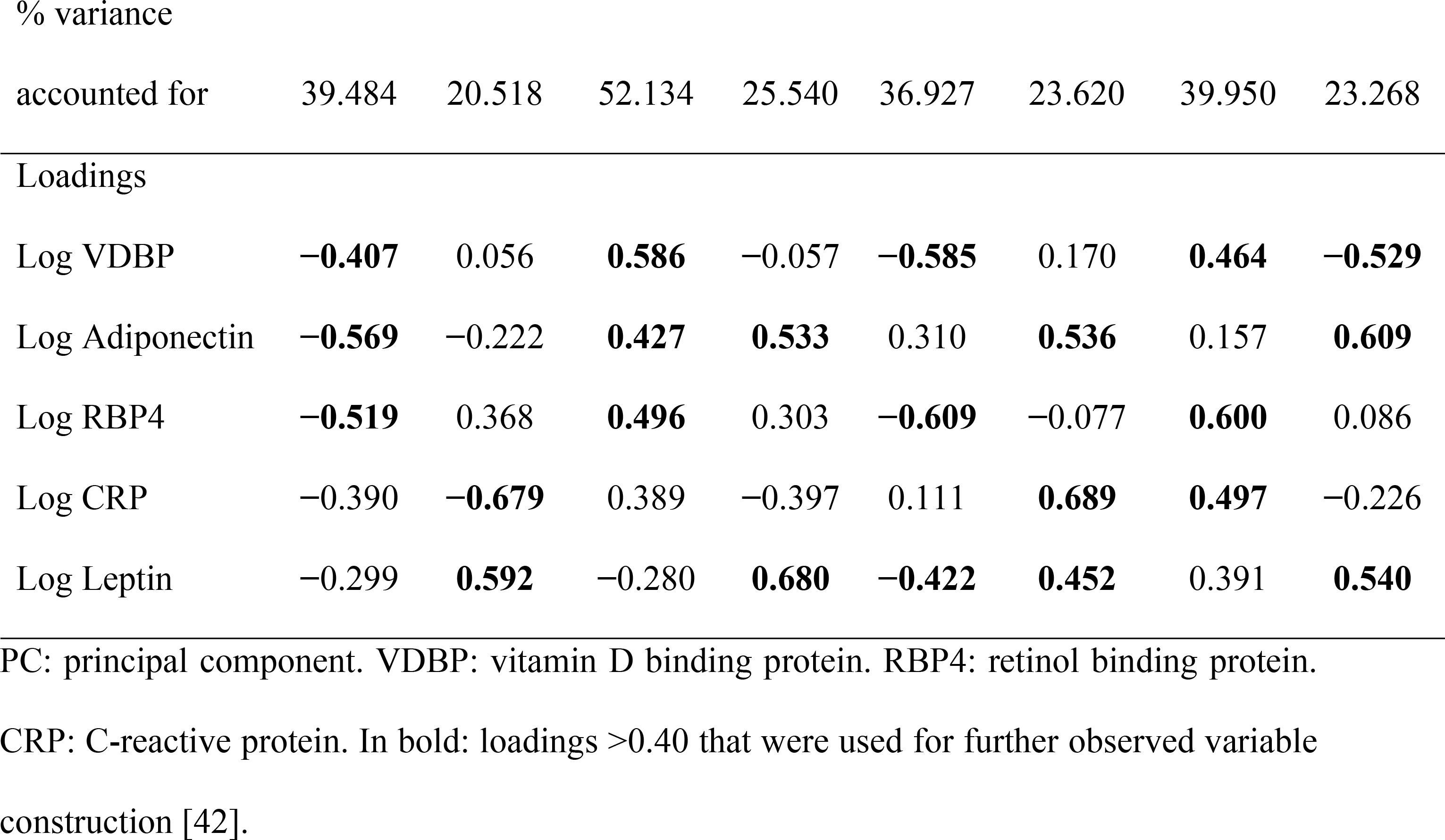

### Association between baseline maternal PC and maternal nutritional status

We analyzed the association between baseline maternal PC and maternal nutrition status including maternal hemoglobin at baseline, maternal height, and maternal MUAC. We found that maternal MUAC was significantly associated with baseline maternal PC1 bp.pc1.D↓A↓R↓ (β = −0.019, *p* = 0.030) (Table 3). Individual maternal biomarkers analysis with maternal nutritional status is presented in S1 Table.

**Table 3.**
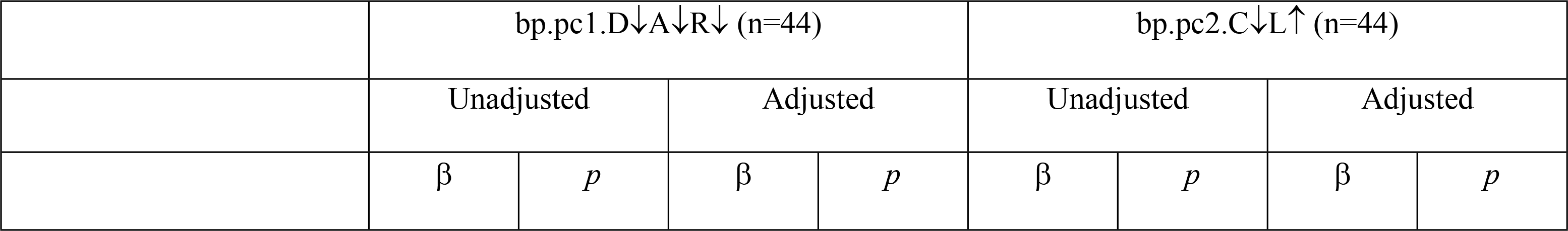
Association between maternal biomarker PC at baseline and maternal nutritional status.

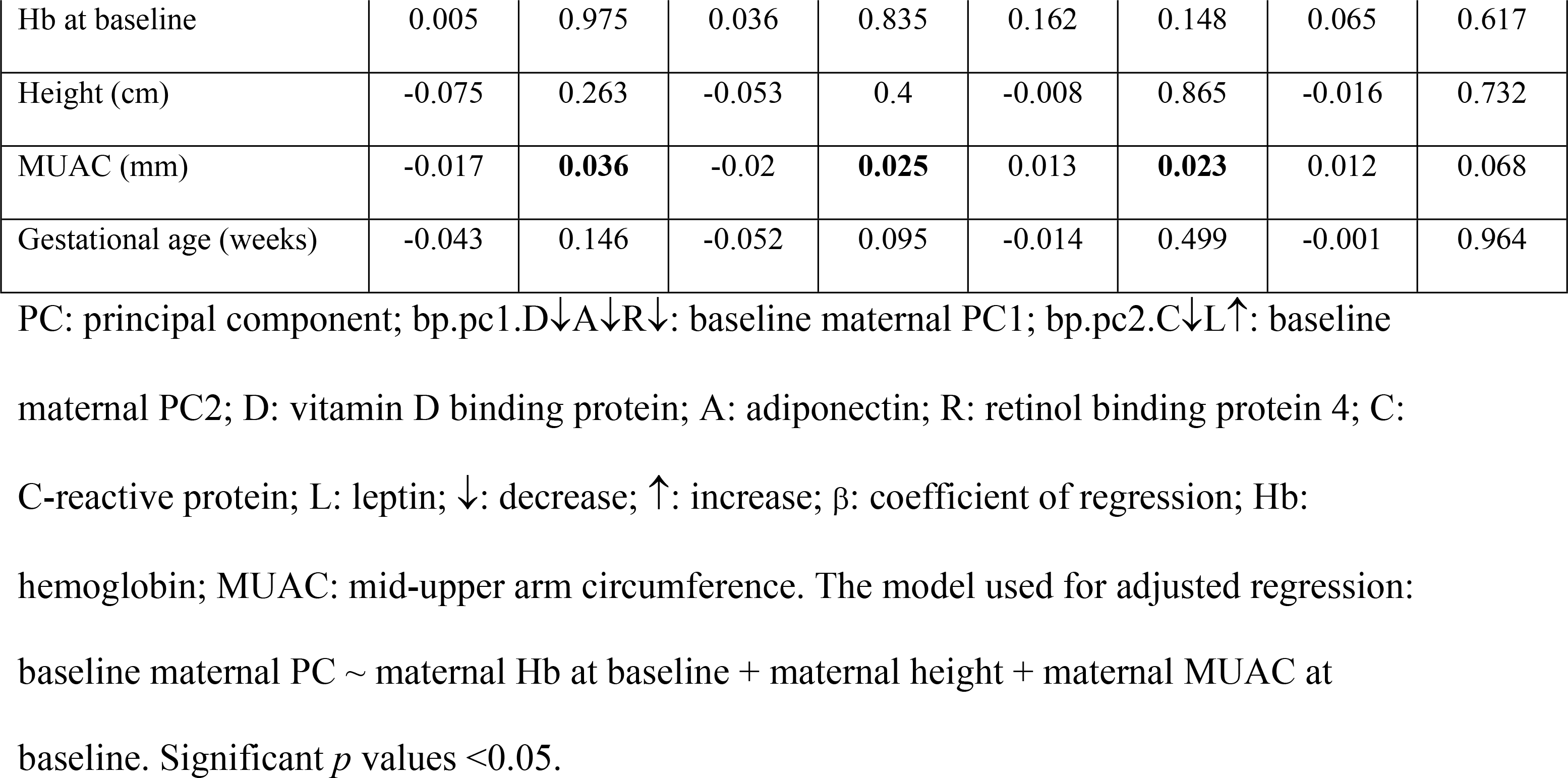

### Association of maternal biomarkers at baseline and post-supplementation

Linear regression results for the associations between maternal biomarker PC scores at baseline and at post-supplementation are presented in Table 4. We found that the baseline maternal PC2 bp.pc2.C↓L↑ was significantly associated with the combined post-supplementation maternal PC1 dp-pp.pc1.D↑↓A↑R↑↓L↓ (β = −0.518, *p* = 0.022). The baseline maternal PC1 bp.pc1.D↓A↓R↓ was significantly associated with the post-supplementation maternal PC2 dp-pp.pc2.A↑C↑L↑ (β = −0.315, *p* = 0.028) (Table 4). Furthermore, we tested the interaction between maternal PC at baseline and supplementation type with maternal PC at post supplementation. We found that the interaction between MMN supplementation and baseline maternal PC2 bp.pc2.C↓L↑ was significantly associated with post-supplementation maternal PC2 dp-pp.pc2.A↑C↑L↑ (*p* = 0.022) (Figure 3A). Individual analysis of maternal baseline and post-supplementation biomarkers is shown in S2 Table.

**Table 4.**
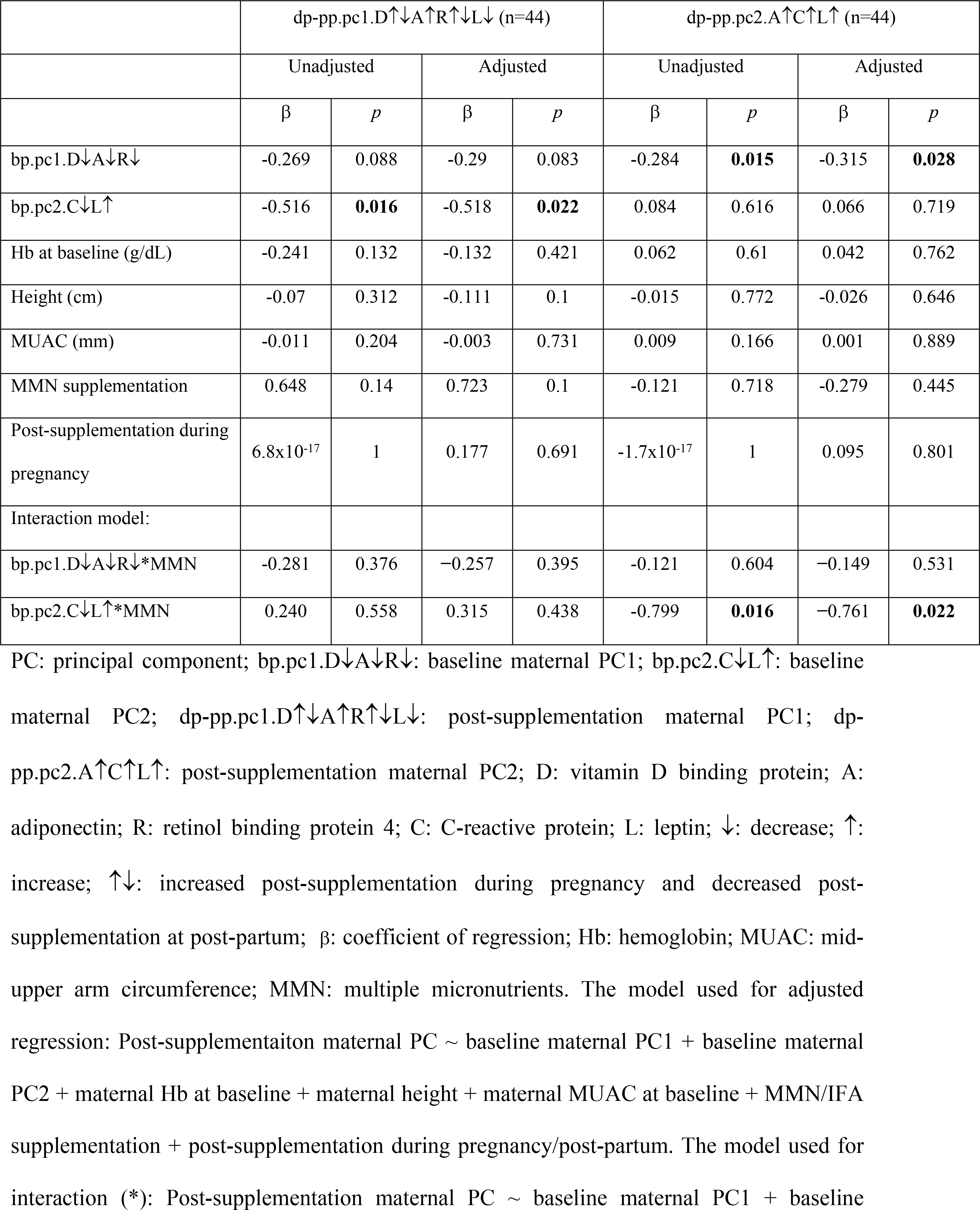
Association between maternal biomarker PC at baseline and post-supplementation.

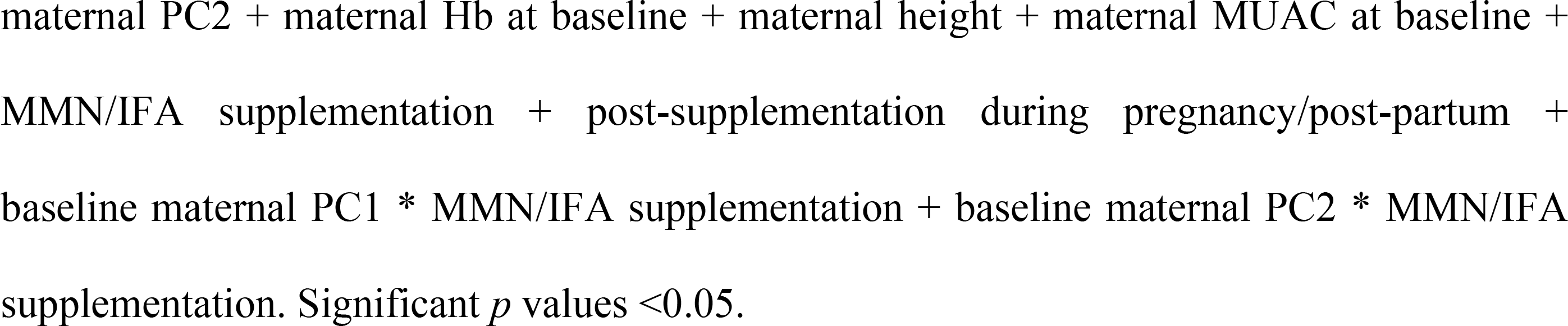

**Fig 3.**
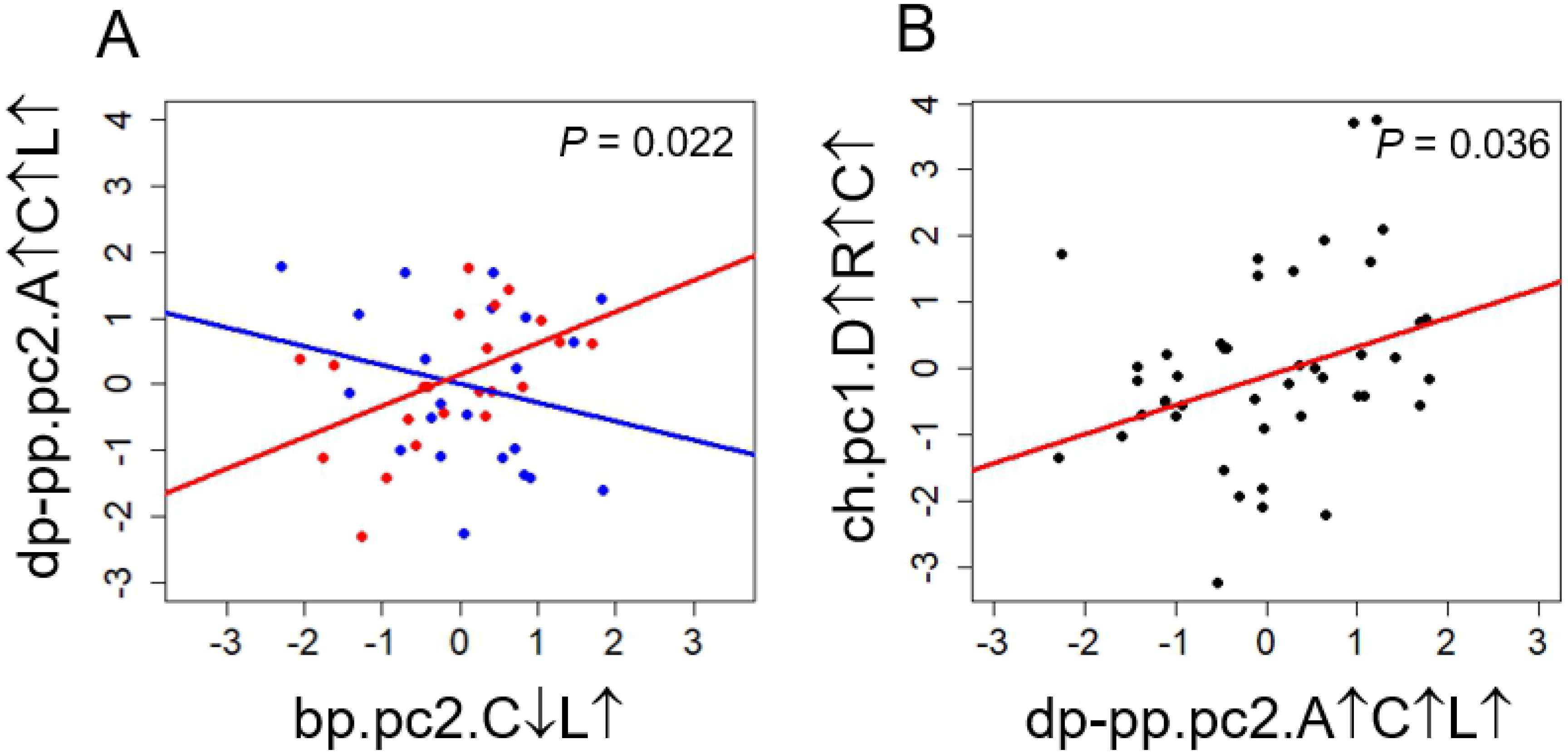
Biomarkers Interaction Plots. **A**. Interaction plot between baseline maternal PC2 bp.pc2.C↓L↑ and supplementation type with post-supplementation maternal PC2 dp-pp.pc2.A↑C↑L↑. Blue line: MMN supplementation; Red line: IFA supplementation. **B**. Interaction plot between maternal PC2 dp-pp.pc2.A↑C↑L↑ and child PC1 ch.pc1.D↑R↑C↑.

### Association of maternal and child biomarkers

We found that the post-supplementation maternal PC2 dp-pp.pc2.A↑C↑L↑ was significantly associated with child PC1 ch.pc1.D↑R↑C↑ (β = 0.439, *p* = 0.036) (Figure 3B). Birth weight was also associated with child PC1 ch.pc1.D↑R↑C↑ (β = −0.826, *p* = 0.036). And we observed that maternal height (β = −0.097, *p* = 0.030), child’s gender boys having lower scores (β = −0.958, *p* = 0.002), and timing of post-supplementation during pregnancy, with samples collected during pregnancy having higher scores (β = 1.224, *p* <0.001, power = 0.972) were significantly associated with child PC2 ch.pc2.D↓A↑L↑ (Table 5). The association of individual child biomarkers with maternal biomarkers at baseline and post-supplementation with child biomarkers are shown in S3 Table and S4 Table.

**Table 5.**
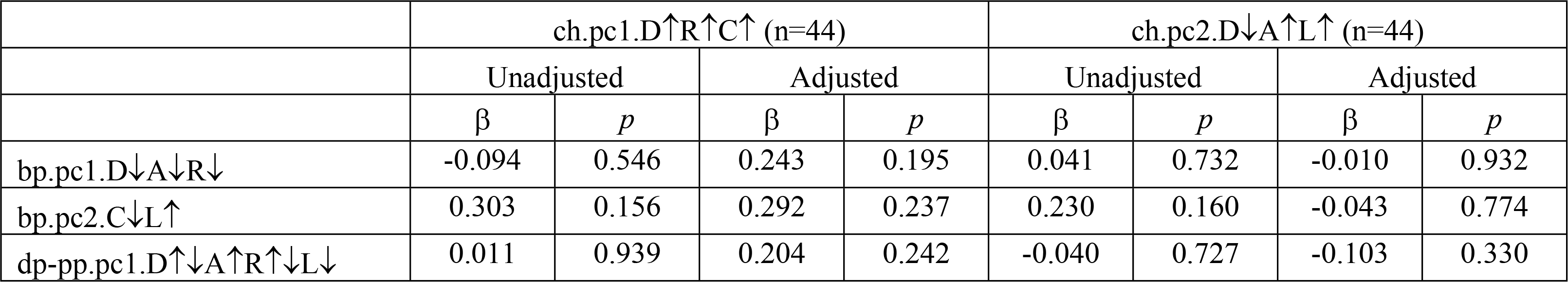
Association between child biomarker PC and maternal biomarker PC.

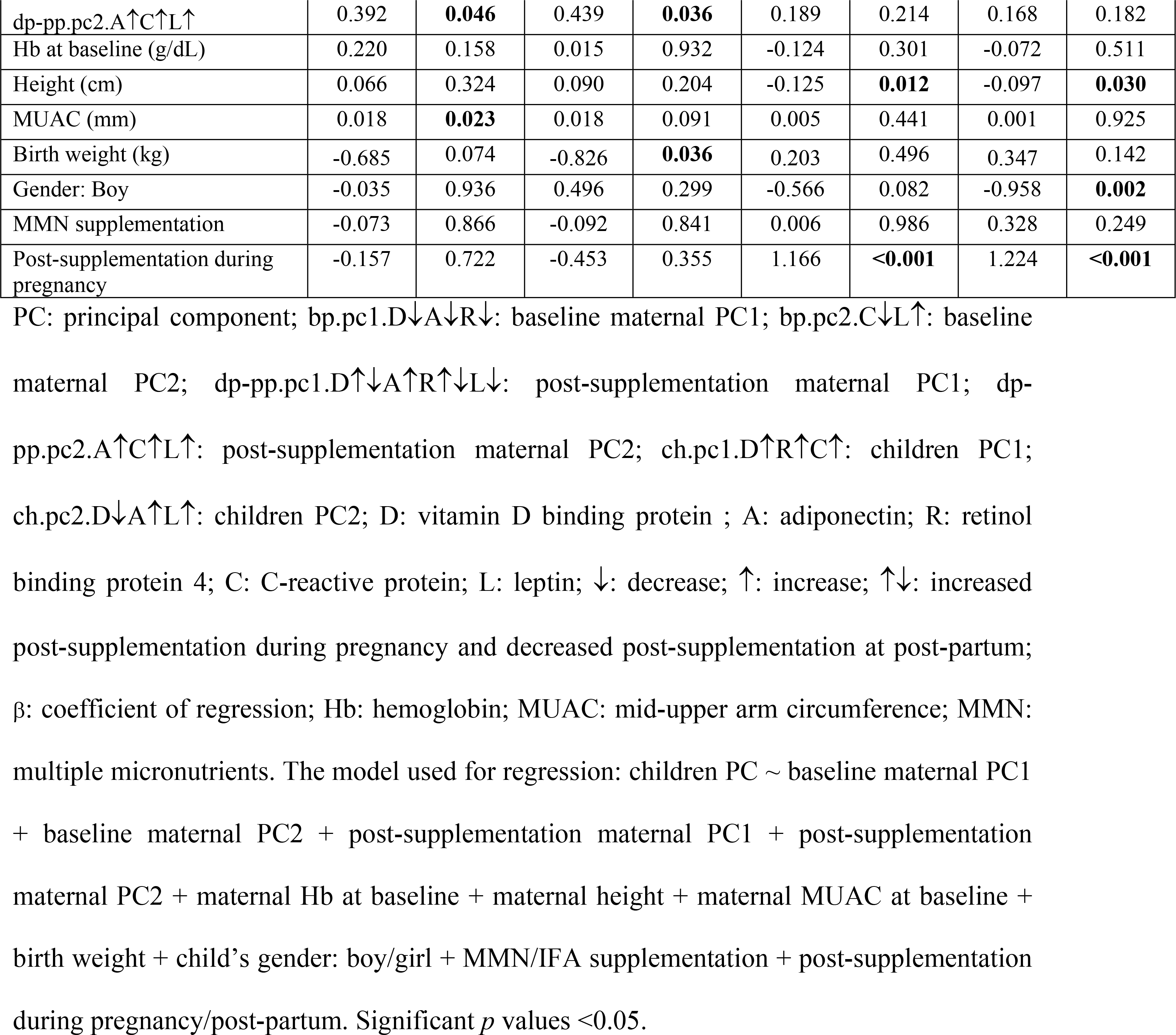

### Association of child health outcome with maternal and child biomarkers

We then analyzed the association of maternal and child biomarker PC scores with child health outcomes (BMIZ, SBP and DBP) with (Table 6). We found maternal dp-pp.pc2.A↑C↑L↑ was negatively associated with child BMIZ (β = –0.302, *p* = 0.022), and maternal dp-pp.pc1.D↑↓A↑R↑↓L↓ (β = 0.224, *p* = 0.036), ch.pc1.D↑R↑C↑ (β = 0.347, *p* = 0.002) and ch.pc2.D↓A↑L↑(β = 0.515, *p* = 0.005) were positively associated with child BMIZ (Figrue 4). Baseline maternal Hb (β = –0.280, *p* = 0.010), maternal MUAC (β = 0.014, *p* = 0.027), and timing of post-supplementation sampling (β = –1.064, *p* = 0.005, power = 0.972) also showed significant association with child BMIZ. No significant associations were found with child SBP and DBP. The association of child health outcome with maternal biomarkers and child biomarkers are shown in S5 Table (maternal biomarkers at baseline), S6 Table (maternal biomarkers at post-supplementation) and S7 Table (child biomarkers).

**Table 6.**
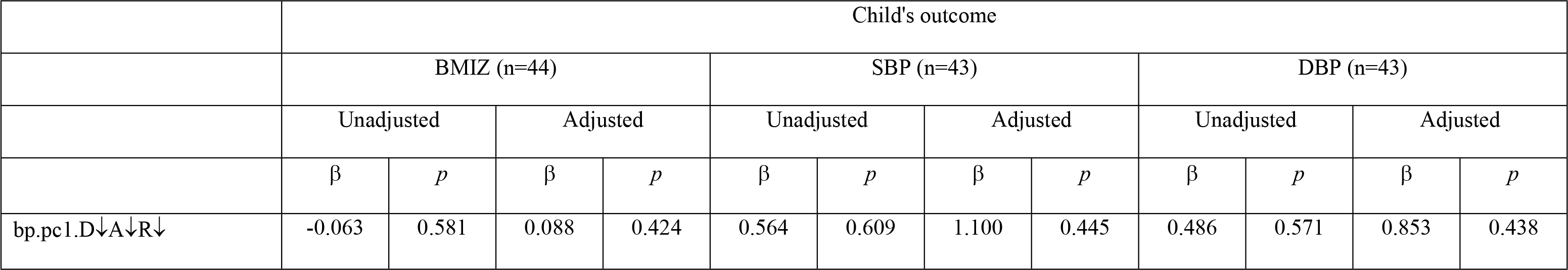
Association between child’s outcome, child’s biomarker PC and maternal biomarker PC.

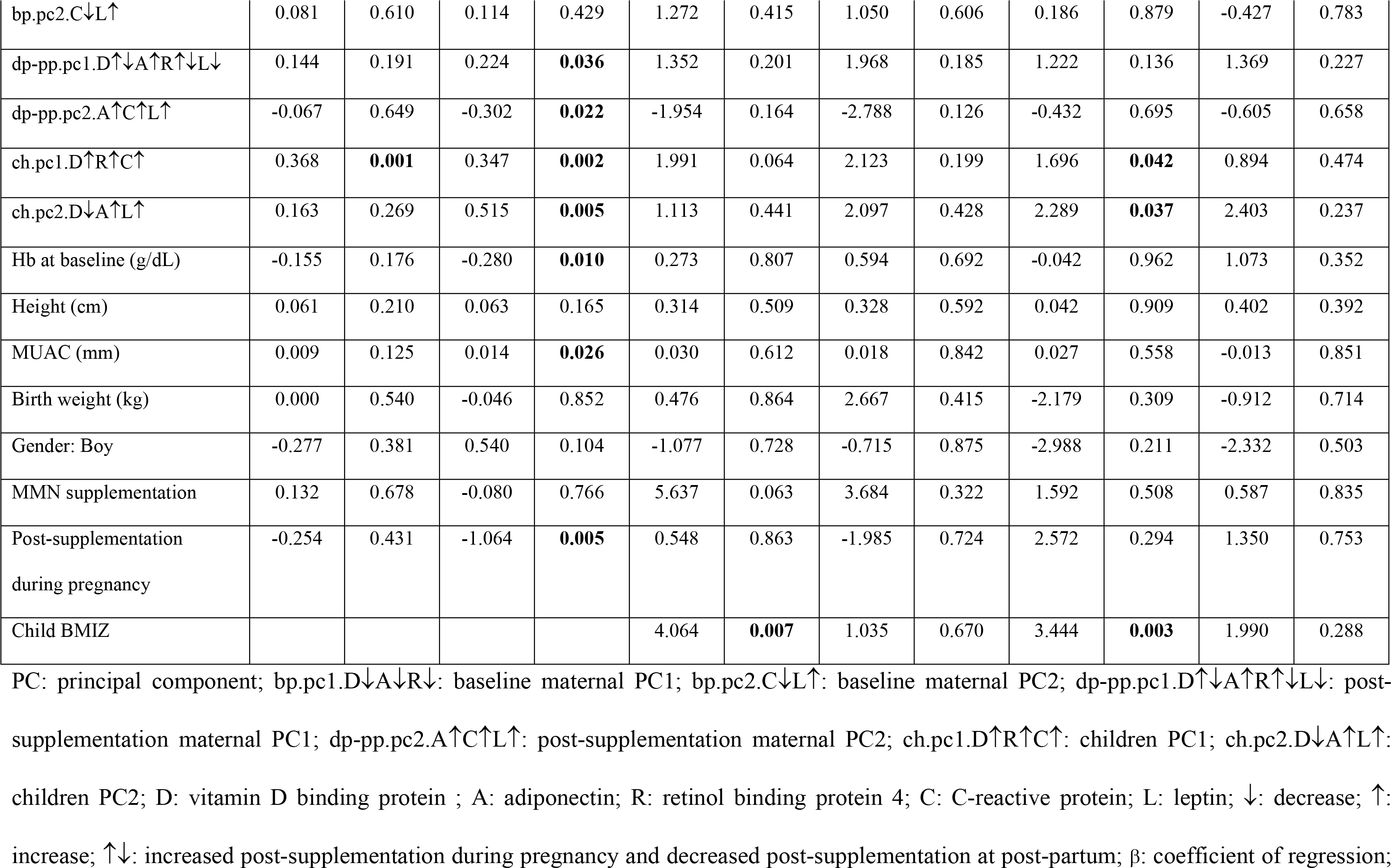

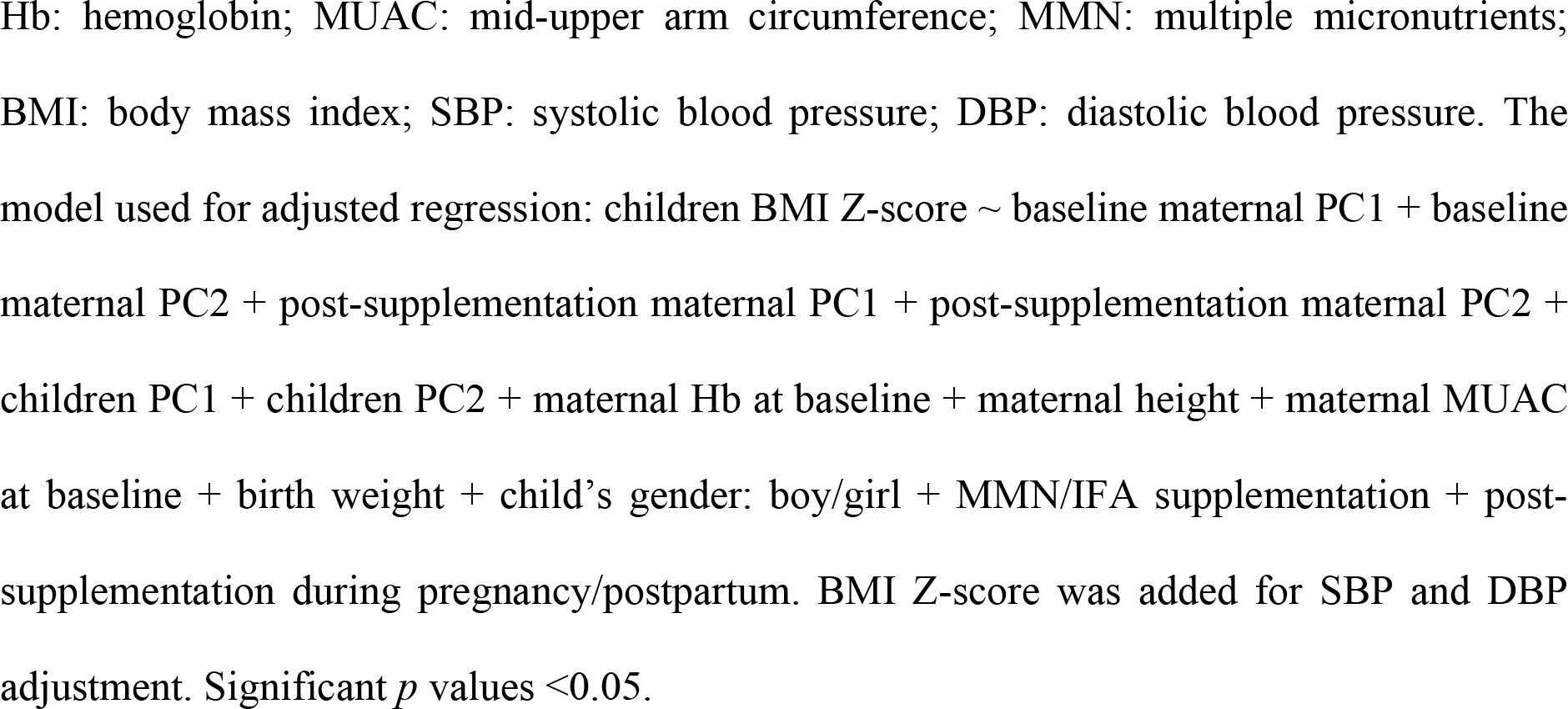

**Fig 4.**
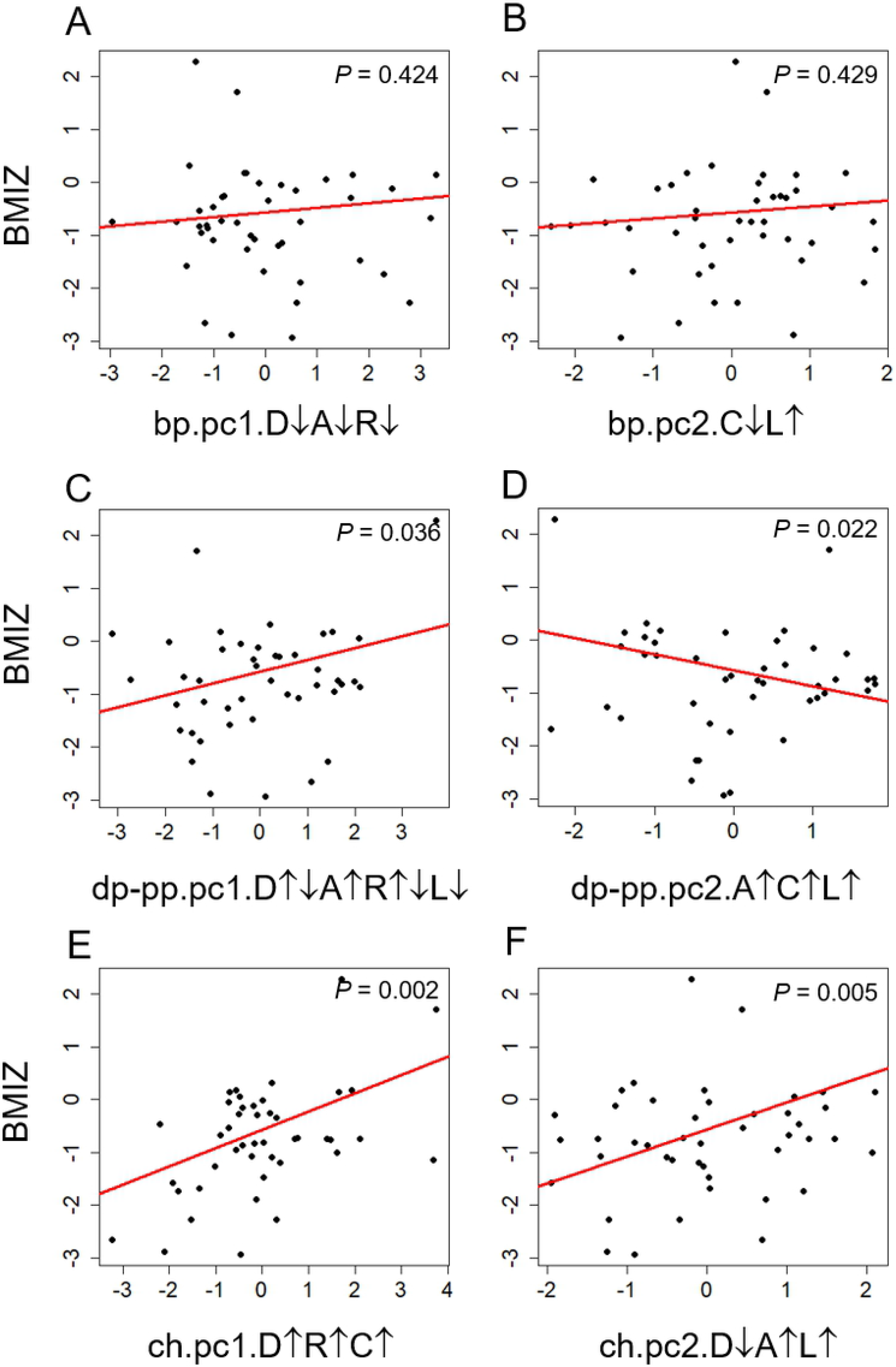
Association of maternal and child biomarkers with BMIZ. **A-B**. Maternal baseline PC1 bp.pc1.D↓A↓R↓ and PC 2 bp.pc2.C↓L↑. **C-D**. Maternal PC 1 dp-pp.pc1.D↑↓A↑R↑↓L↓ and PC 2 dp-pp.pc2. **E-F**. Child ch.pc1.D↑R↑C↑ and PC 2 ch.pc2.D↓A↑L↑.

## DISCUSSION

To our knowledge, few studies have explored the association of maternal metabolic biomarkers during pregnancy and post-partum with child metabolic biomarkers at age 9-12 years. Moreover, because biomarkers may not work independently, but in concert, potential interactions between patterns of biomarker components may better represent the complexity of their downstream effects. Therefore, rather than analyze the effects of individual biomarkers, we utilized PCA to identify patterns that represented their covariance structure, and analyzed the associations between the resulting PC and other PC and outcomes.

PCA showed that maternal biomarkers at baseline and post-supplementation during pregnancy and post-partum had distinctive component structures, indicating that gestational age influences the maternal biomarker patterns. We found that maternal MUAC was associated with baseline maternal PC, indicating the influence of maternal nutritional status on maternal biomarkers. This notion has been previously reported where nutritional status measured by BMI was positively correlated with leptin, adiponectin, and RBP4 concentrations [43–45], though these studies were not done in pregnant women.

We also found that maternal biomarkers PC at baseline were associated with biomarkers PC at post-supplementation, although cross associations in these timepoints between individual biomarkers were observed only in adiponectin and RBP4 (S2 Table). This indicates that biomarkers are working together to influence each other in the biological system. Moreover, MMN supplementation interacted with maternal biomarkers PC at baseline to influence biomarkers PC at post-supplementation. Previous studies have reported that vitamin C and E supplementation reduced CRP concentrations [36,46], and vitamin D influenced serum leptin and adiponectin concentrations [47].

We observed that maternal PC scores at post-supplementation were positively associated with child PC scores at 9-12 years. Previous studies showed that maternal leptin concentration was correlated with children’s leptin concentration in cord blood [23,48] and serum of 9-years old children [49]. Postpartum maternal biomarkers may be associated with child biomarkers through breast milk, in agreement with a previous study that reported a correlation between leptin concentration in breast milk with its concentration in maternal serum and infant’s weight gain [50]. Although genetics was also reported to have moderate influences for the variation of biomarkers concentration [51,52], environmental factors such as nutrition, including micronutrients, and infection have been reported to also modulate adipocytokines and inflammatory markers concentrations [32–37,53]. Our analysis did not include the influence of dietary intake on biomarkers concentrations, which could reveal additional associations.

BMI-for-age z-score represents nutritional and health conditions in children and adolescents [54]. Our study showed that maternal and child biomarker PC scores were positively associated with child BMIZ. This is in line with previous studies that reported BMI in children was correlated with biomarkers concentration, such as leptin [55] and RBP4 concentrations [56]. BMIZ in children was reported to be influenced by in-utero milieu, such as smoking during pregnancy that was associated with lower BMIZ [57]. In our study, the average BMIZ was below WHO standard [41], which means the children tended to be underweight. However, BMI is modifiable, and can be improved by nutritional and behavior interventions [58]. Thus, maternal MMN supplementation during pregnancy might indirectly influence child BMIZ considering that our results indicated that MMN modified the association between maternal baseline and maternal post-supplementation biomarkers, while maternal post-supplementation biomarkers are associated with both child biomarkers and BMIZ.

It has been suggested that pre-pregnancy and pregnancy nutritional status have long term effects on health outcomes of children. Maternal height was positively associated with child PC scores, and maternal MUAC also had a positive association with child PC scores, although not significant. Maternal Hb during pregnancy and height were also associated with children’s BMIZ. These results support the potential influence of maternal nutritional status on the children metabolism and health. This notion has been previously reported where maternal nutritional status measured by BMI was correlated with children’s nutritional status measured by BMI [59] and weight for height z-score (WHZ) [60]. Maternal BMI was also reported to be associated with infant serum leptin values [45]. Therefore, our finding again emphasized the importance of optimal macronutrients intake during pregnancy that would improve maternal nutritional status and later child’s health [61].

We proposed that maternal biomarkers of adipocytokines and inflammatory markers could influence the same biomarkers in the child through the interactions of immunologic and metabolic factors. Adiponectin, RBP4, CRP, and leptin play important roles in regulating metabolism, energy homeostasis, and inflammatory responses, while VDBP has a role in modulating immune and inflammatory response. The immune and metabolic system have co-evolved to signal each other and form complex networks as a response to environmental exposures, such as the secretion of leptin and adiponectin that are contra-regulated [62,63]. Transfer of immune and metabolic properties between mother and her child occurs through the placenta during pregnancy [23,64], and through breast milk during the neonatal period [50]. Together, these immune-metabolic signals provide innate and adaptive immunities and influence the metabolic homeostasis of the newborn. The transmission of these cross-generational immune and metabolic properties may be modified via optimal macronutrients and micronutrients intake during pregnancy and postpartum. Maternal adverse conditions, such as malnutrition or infection may modify these signals and alter the newborn immunity, consequently influencing newborn and infant health, and possibly later life [65,66].

Despite the sparsity of the data, which is the limitation of this study, the consistency of the associations observed is of interest. Nevertheless, replication of this study’s findings would be warranted. In addition, due the multiple hypotheses tested, the multiple comparisons in the study were unavoidable, but again we note the consistency of the findings. To our knowledge, this is the first study with complete mother-child dyads showing the effect of MMN on the child outcomes via changes of the mother’s biomarkers. As with other effect of nutrients, the MMN effect cannot be determined based on a single biomarker only, as there would be many pathways involved. Therefore, analyzing the effect of a particular pattern of combined relevant biomarkers is more relevant, as conducted here.

In conclusion, maternal biomarkers of adipocytokines and inflammatory markers PC during pregnancy were modulated by MMN supplementation, and associated with child PC scores of the same biomarkers 9-12 years later, as well as with children’s BMIZ. Improving maternal nutritional status may improve child’s conditions not only directly after birth, but also during their childhood, and until adulthood.

## ACKNOWLEDGEMENTS

We thank all the pregnant women, their children, the families, the communities, and the midwives in Lombok, West Nusa Tenggara, who participated in and facilitated the original SUMMIT and its follow up studies. We are grateful to Dr. Husni Muadz of Mataram University, Center for Research on Language and Culture, and Ms. Mandri Apriatni, SPd, MA, of Summit Institute of Development, who were involved in the SUMMIT study, and Mr. Miswar Fattah and his team of Prodia CRO for the use of their MagPix CCD Imager. The authors are grateful to the Scientific Members of the Indonesian Danone Institute Foundation and Dr. Jacques Bindels of the Asia Pacific R&D Danone Baby Nutrition, for their inputs during the initial development of this study.

